# A Mathematical Model of Dietary Lipid Absorption and Postprandial Chylomicron Dynamics

**DOI:** 10.64898/2026.06.25.734455

**Authors:** Christian Simonsson, Oscar Silfvergren, Henrik Podéus, Kajsa Tunedal, William Lövfors, Karin G. Stenkula, Elin Nyman, Gunnar Cedersund

## Abstract

Obesity and related conditions such as dyslipidemia impose an increasing burden on healthcare systems worldwide. These conditions are associated with altered postprandial chylomicron (CM) metabolism, the elusive and critical first step in lipid metabolism. This step remains elusive because it is governed by large interindividual variations and a complex set of intestinal processes. In particular, the second meal effect (SME) implies that enterocytes release previously stored fat during subsequent meals. To deal with this complexity, CM and lipid metabolism have previously been explored using mathematical modeling. However, existing models primarily describe TAG dynamics following a single meal or are too complex for practical personalization across datasets. Herein, we address these limitations by presenting a small-scale mathematical model of CM dynamics that incorporates the SME. The presented model successfully describes data from six clinical studies of both single and repeated meal interventions. Model performance was further evaluated by predicting independent datasets using a BMI-dependent calibration. Finally, to demonstrate model applicability, we simulated full-day responses consisting of three sequential meals in individuals with varying BMI values, with qualitative agreement to clinical observations. This work supports our understanding of the SME, person-specific CM postprandial responses, and mechanisms underlying obesity.

## Introduction

Obesity and related metabolic conditions, such as insulin resistance, atherosclerosis, and type 2 diabetes (T2D), are placing an increasing burden on our healthcare systems. In T2D research, mathematical models of insulin and glucose regulation have substantially advanced our understanding of disease progression and helped facilitate the development of treatment strategies (1,2). In contrast, fewer mathematical models have focused on postprandial lipid metabolism. Dyslipidemia, characterized by elevated circulating lipid levels, is an important feature of metabolic disease progression and may in part be driven by the postprandial appearance of chylomicrons (CM) (3). As CM dynamics are highly complex, incompletely understood, and subject to substantial inter-individual variation, the development of robust mathematical models of CM appearance and postprandial metabolism may be important for advancing our understanding and treatment of metabolic diseases.

A mathematical model of postprandial CM dynamics needs to describe key biological mechanisms. One such mechanism is the intestines’ ability to store triglycerides (TAG) for later release, which contributes to a lipid absorption phenomenon known as the ‘second meal effect’ (SME) (4,5). The SME describes how subsequent meals will result in higher CM-TAG release compared to the first meal, even when the meals are identical (6). Several studies have observed the SME in humans (4,7–11), demonstrating complex relationships between CM-TAG release, prior meal composition, and the ingestion of other macronutrients. For example, Evans et al. showed that a low-fat meal, after a high fat breakfast, still triggered a CM-TAG release (4). Moreover, CM-TAG appearance differs between individuals, with reported differences related to sex, insulin sensitivity, and lipid-form (emulsion or solid) (12,13).

Only a handful of mathematical models have been developed to describe CM-TAG release. One example of such a model is the model by Leohr et al. (14), a semi-physiological model that was used to quantify the CM-TAG response to a high-fat meal in both obese and healthy individuals. While this model describes CM-TAG data for a single meal, it does not aim to capture the SME. Other models describe TAG dynamics after a single meal (15), but these do not explain CM-TAG dynamics. Sosa et al. (11) focused primarily on CM release and investigated differences in CM appearance and the SME between insulin-sensitive and insulin-insensitive individuals, through both experimental data collection and systems biology analysis. Their analysis focused on a single study of sequential meals, and employed a mathematical model, supporting the hypothesis that stored enteral TAG bypasses re-synthesis and lipolysis, enabling rapid secretion. However, there is still a need for a simple but physiologically meaningful model, integrating data from different studies, capable of reproducing the complex dynamics of the SME, while remaining suitable for individualized parameterization.

Herein, we present such a mathematical model designed to describe CM plasma appearance from repeated mixed meals of several independent studies (**Fig. 1A**), supporting our understanding of the SME and mechanisms underlying obesity.

**Figure 1.**
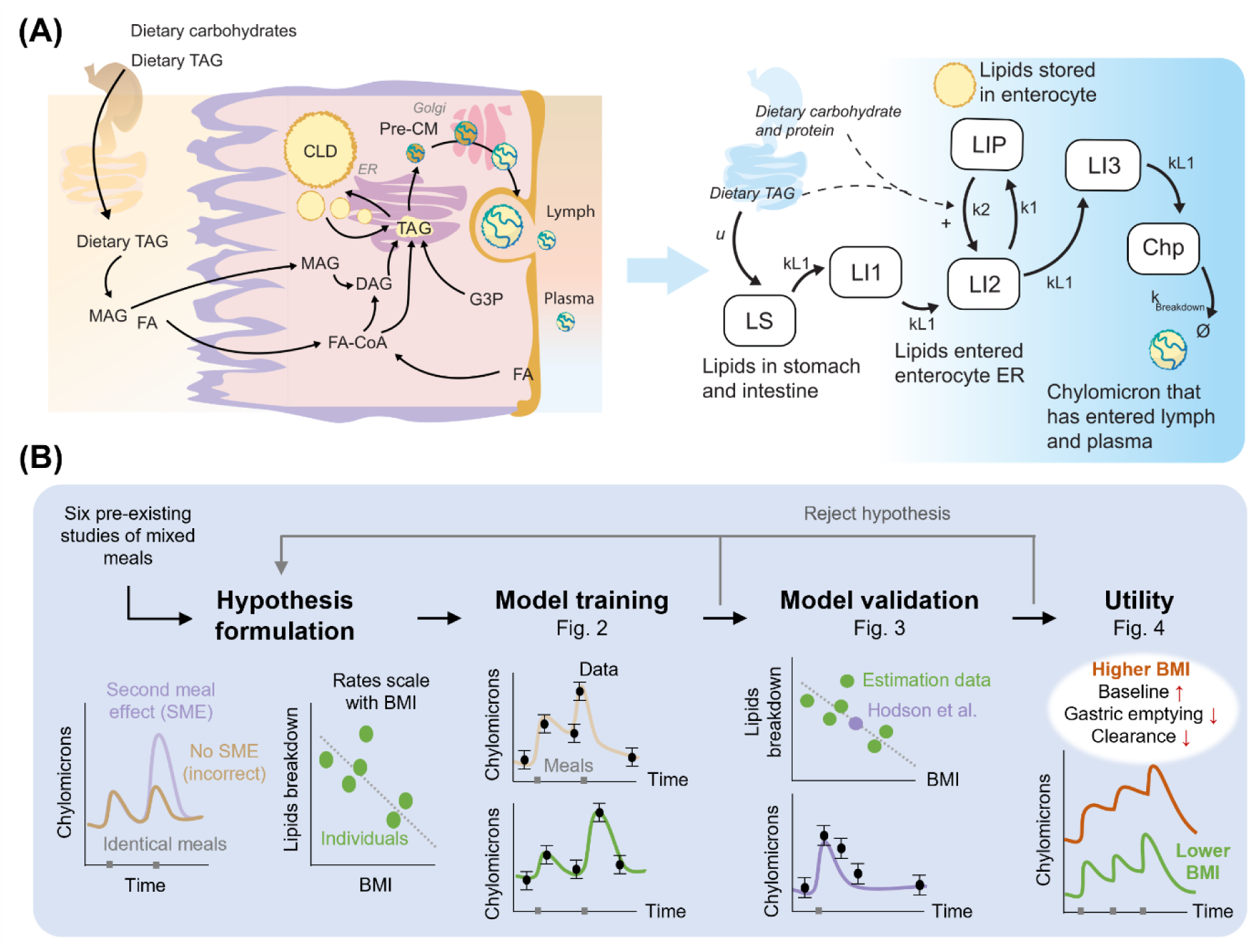
Study setup and approach. **(A)** Physiological processes involved in enterocyte lipid absorption from intestine (left) translated into a mathematical model (right). **(B)** Methodology and workflow. The model development was conducted in stages: Hypothesis formulation, where a hypothesis was translated into a mathematical model; Model training, where the model was trained to six heterogenous studies both individually (**Fig. S1**) and jointly; Model validation, where the model’s ability to predict independent data was tested; and Utility, where the model was used to predict various repeated meal interventions for individuals with different BMI.

## Results

In this study, we have iteratively developed and evaluated a mathematical model for plasma CM-TAG dynamics following both single and multiple meals (**Fig. 1A**). As a first step, the model was trained individually to six different studies (**Fig. S1**), where two model parameters exhibited substantial interpopulation variability (**Figs. S2–S3**). As a second step, the mathematical model was trained to all studies jointly while allowing group-specific variability (study and intervention) in the two parameters identified with substantial variability (**Fig. 1B**, model training). Both parameters were found to be negatively associated with the mean body mass index (BMI) reported in the corresponding studies, using linear regression. As a third step, this relation between BMI and model parameters was used to predict CM dynamics of an independent dataset not used in model training (**Fig. 1B**, model validation). Finally, the established model was used to investigate repeated sequential meals in individuals with varying BMI values (**Fig. 1B**, utility).

### Individual model agreement with CM dynamics for single and multiple meals

The mathematical model of CM postprandial dynamics can describe six different studies individually (**Fig. S1**). Details regarding each intervention and cohort in the training dataset can be found in the supplementary material (**Table S1**). The agreement between model simulations and data was evaluated using χ^2^-tests with 95% confidence interval (see material and methods). The model passed the χ^2^-test for all studies individually (**Table 1**).

**Table 1.** Model agreement when fitted to all data sets individually, with corresponding χ^2^-test.

| Study | $\chi^2$-test |
| --- | --- |
| Pramfalk *et al.*, (12) males (**Fig. S1A**) | $0.2 < T_{\chi^2} = 21$ |
| Pramfalk *et al.*, (12) females (**Fig. S1B**) | $0.1 < T_{\chi^2} = 21$ |
| Milan *et al.*, (16) young (**Fig. S1C**) | $0.1 < T_{\chi^2} = 12.6$ |
| Milan *et al.*, (16) old (**Fig. S1D**) | $0.3 < T_{\chi^2} = 12.6$ |
| Evans *et al.*, (4) water (**Fig. S1E**) | $9.1 < T_{\chi^2} = 18.3$ |
| Evans *et al.*, (4) low-fat (**Fig. S1E**) | $6.0 < T_{\chi^2} = 18.3$ |
| Evans *et al.*, (4) water (**Fig. S1F**) | $9.1 < T_{\chi^2} = 18.3$ |
| Heath *et al.*, (17) healthy (**Fig. S1G**) | $15.4 < T_{\chi^2} = 18.3$ |
| Vors *et al.*, (13) normal – lipid spread (**Fig. S1H**) | $3.3 < T_{\chi^2} = 17.0$ |
| Vors *et al.*, (13) normal – lipid emulsion (**Fig. S1I**) | $3.7 < T_{\chi^2} = 17.0$ |
| Vors *et al.*, (13) obese – lipid spread (**Fig. S1J**) | $0.52 < T_{\chi^2} = 17.0$ |
| Vors *et al.*, (13) obese – lipid emulsion (**Fig. S1K**) | $2.7 < T_{\chi^2} = 17.0$ |
| Barrows *et al.*, (9) healthy (**Fig. S1L**) | $2.6 < T_{\chi^2} = 31.4$ |

A parameter analysis was performed to evaluate the interstudy differences (**Figs. S2-S3**). We found that two parameters displayed substantial interstudy variability: *kBreakdown* (governing the rate of CM clearance), and *kL1* (governing the rate of transport in intestine and enterocytes). This assessment was performed by comparing the parameter boxplot distributions (**Fig. S2**) and parameter-to-parameter correlations (**Fig. S3**). For the boxplot distributions, the confidence intervals (CIs) for both *kL1* and *kBreakdown* appeared broader for intervention including single meals, in comparison to intervention including multiple meals. For the parameter correlations, the analysis indicates that *kL1* and *kBreakdown* correlate, and thus both could potentially be described with a joint description. Together, these insights indicate that intervention and study variability could be described by allowing freedom in these two identified parameters.

By allowing freedom in the parameters *kBreakdown* and *kL1,* the model can simultaneously describe all estimation data (**Fig. 2**), confirmed by a χ^2^-test with 95% confidence interval (90.1*<T_χ2_*=148.8). Additional tests were performed to evaluate if only one parameter was sufficient to represent interstudy differences (**Fig. S4,** freedom only in *kBreakdown;* **Fig. S5,** freedom only in *kL1*). We found that freedom in only one of these parameters could pass the χ^2^-test (*kBreakdown*, 144 *< T_χ2_*=148.8; and *kL1,* 132.4 *< T_χ2_*=148.8). However, both cases were not accepted, as they did not sufficiently capture the SME and qualitative assessment of the meal responses (**Figs. S4-S5**). Therefore, to more accurately represent interstudy variability, model development proceeded with allowed freedom in both parameters (**Fig. 2**).

**Figure 2.**
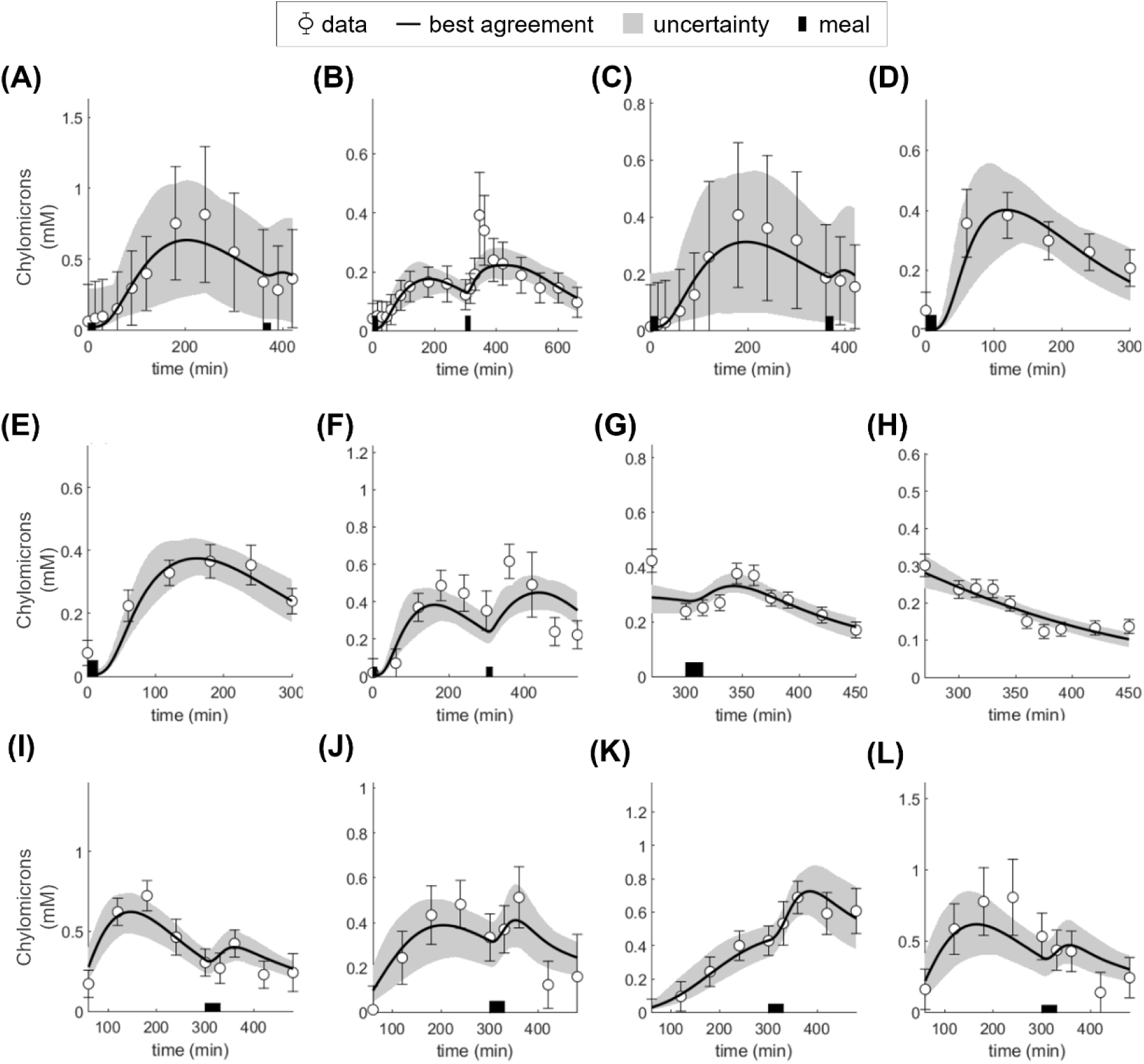
Model simultaneous agreement to all estimation data. For all panels: area indicates model uncertainty, errorbars represent data, lines represent model best agreement with data, and thick black bar(s) on x-axis indicate meal consumption. **(A)** Pramfalk et al., (12) males **(B)** Barrows et al., (9) healthy. **(C)** Pramfalk et al., (12) females. **(D)** Milan et al., (16) young, **(E)** Milan et al., (16) old. **(F)** Heath et al. (17) healthy. **(G)** Evans et al. (4) low-fat. **(H)** Evans et al. (4) water. **(I)**Vors et al. (13) normal spread. **(J)** Vors et al. (13) normal emulsion. **(K)** Vors et al. (13) obese spread. **(L)** Vors et al. (13) obese emulsion.

### Using the model to predict independent validation data of CM appearance

Values of the two parameters describing study- and population-differences (*kBreakdown* and *kL1*) have to be assumed to be able to make predictions of new individuals or meal interventions (**Fig. 3**). To address this, the relationship between the study-specific parameters and known study covariates, e.g., *BMI,* age, and *Homeostatic Model Assessment for Insulin Resistance* (HOMA-IR) (**Table S1**), was assessed via a linear-regression analysis (**Fig. 3A-B**). This was performed using the parameter-value (in log_e_) corresponding to the best model agreement with data. We found that the highest correlation was observed between BMI and *kBreakdown* with a Pearson correlation coefficient of R = 0.63 (**Fig. 3A**). A slightly smaller correlation coefficient was observed between BMI and *kL1 (R = 0.57) (****Fig. 3B***). Using these regression equations, we could now get a plausible value of the calibration parameters based on the known BMI of an individual.

**Figure 3.**
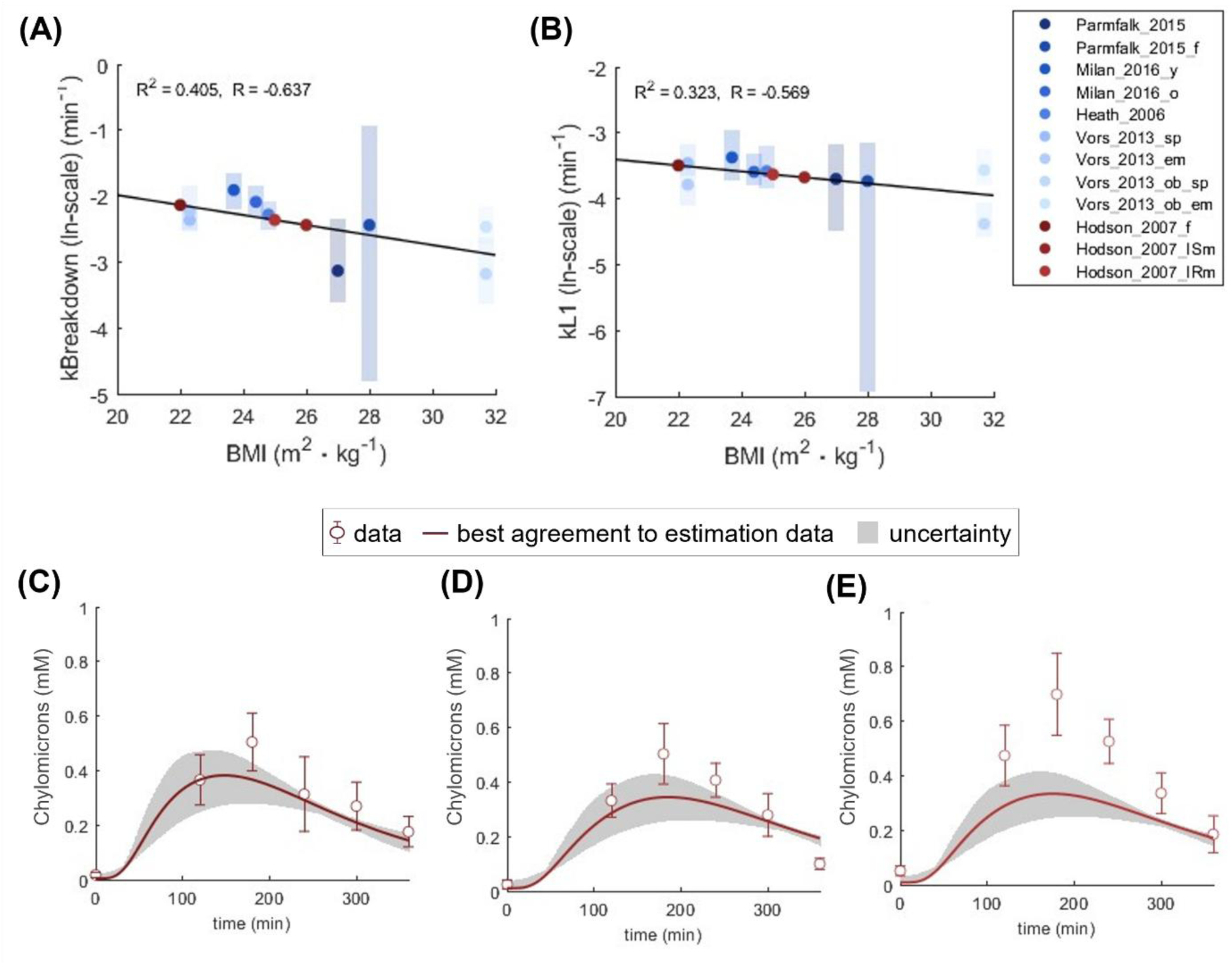
Model predictions of interventions from an independent study (Hodson et al.) (18). For all panels: area indicates model uncertainty, and errorbars represent data. **(A)** The value of kBreakdown for the new cohorts was estimated using the regression model based on BMI. Dots show best found agreement with data. The line shows trends using linear regression. **(B)** A corresponding regression model was used to estimate the kL1 values. Dots show best found agreement with data. The line shows trends using linear regression. **(C)** The model prediction of the cohort comprised of healthy women from Hodson et al. (18). Errorbars show population-data. **(D)** The model prediction of the insulin sensitive men cohort from Hodson et al. (18). Errorbars show population-data. **(E)** The model prediction of the insulin resistant men cohort from Hodson et al. (18). Errorbars show population-data.

We used the BMI-dependent parameter calibration to make predictions of an independent study (18) (**Fig. 3C-E**). In this study, plasma CM was measured after a single meal in three different cohorts: women (**Fig. 3C**; mean BMI of 22), insulin sensitive men (**Fig. 3D**; mean BMI of 26), and insulin resistant men (**Fig. 3E**; mean BMI of 25). The model simulations agreed with the data from the women cohort (**Fig. 3C**) and the data from the cohort of insulin sensitive men (**Fig. 3D**), whereas the model underpredicted the CM for the insulin resistant men (**Fig. 3E**).

### Model predictions of daily CM variations for different individuals and meal patterns

To highlight the usability of the constructed model, we investigated CM appearance during three sequential meals (breakfast, lunch, and dinner containing 15 g, 30 g, and 30 g of fat respectively) during a day (**Fig. 4**). The simulations were performed using different BMI inputs to visualize the relation between CM dynamics and metabolic dysregulation (**Fig. 4A**). Here, we can observe that increasing BMI (used herein as a proxy for metabolic dysregulation) is connected to higher peaks of CM during the day. To highlight this behavior further, we simulated separate CM dynamics for individuals with different BMI (**Fig. 4B**) and mean levels of circulating CM (**Fig. 4C**). Together, these three figures (**Fig. 4A-C**) indicate increasing CM peak value with higher BMI and higher mean CM throughout the day. More specifically, the model predicts postprandial chylomicron exposure approximately 2-fold higher for people with a BMI of 30 compared to a BMI of 20. This is in qualitative agreement with literature, where obese individuals (BMI 30 or above) have been observed to have a 2- to 3-fold higher postprandial chylomicron exposure than lean participants (19). Similar observations have been reported in other works (20,21).

**Figure 4.**
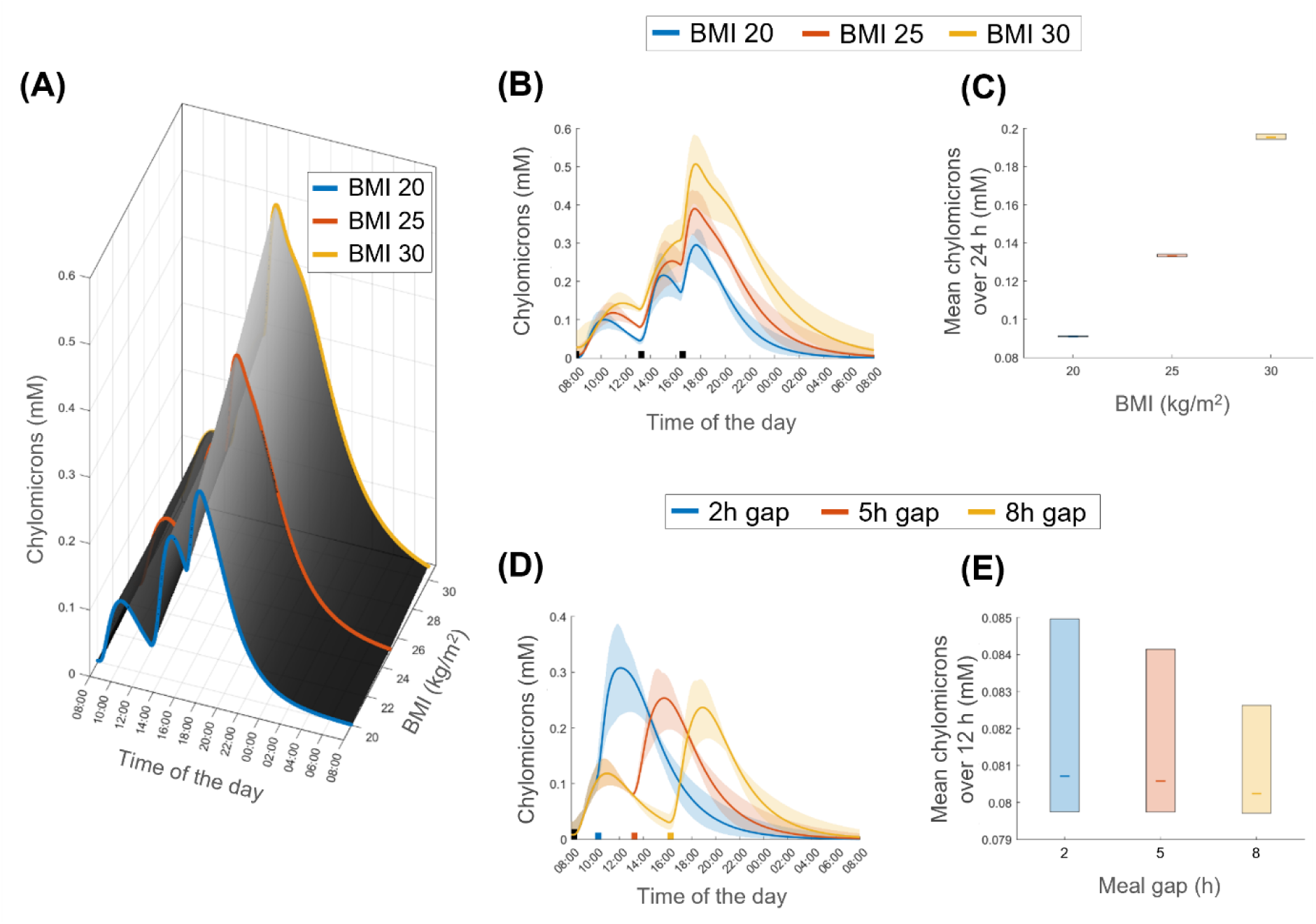
Model prediction of one day consisting of three meals of individuals with different BMI and time between meals. **(A)** Relationship between plasma chylomicron dynamics and BMI over a day of eating three meals. The colored black area is predicted plasma CM, and colored lines show three BMI profiles: BMI of 20 (blue), BMI of 25 (orange), and BMI of 30 (yellow). **(B)** Simulations of chylomicron dynamics for three different BMI groups: BMI of 20 (blue), BMI of 25 (orange) and BMI of 30 (yellow). The meals consisted of 15, 30, and 30 g lipids respectively. **(C)** Predicted mean chylomicron values during the 24 hours shown in Figure 4B. Lines represent the mean and colored boxes represent model uncertainty. **(D)** Simulations of chylomicron dynamics for three different groups with different meal timing; 2-hours gap from first meal (blue), 5-hours gap (orange), and 8-hours gap (yellow). The first meal consisted of 15 g and the second of 30 g of lipids **(E)** Predicted mean chylomicron values during the 24-hours shown in Figure 4D. Lines represent the mean and colored boxes represent model uncertainty.

We also simulated how meal patterns affect the mean level of CM during the day (**Fig. 4D-E**). The model predicted that increasing the time between meals reduces the SME, with a substantial reduction between having a 2-hours gap and a 5-hours gap between meals (**Fig. 4D**, blue compared to orange). Importantly, we found that mean levels of circulating CM also decreased with longer time periods between meals (**Fig. 4E**). In this meal-timing investigation, a SME is predicted when a meal is consumed 5-hours post the first meal (**Fig. 4D**, orange), which is in agreement with clinical observations (22). Furthermore, the model predicts a diminishing SME effect between 5-hours and 8-hours (**Fig. 4D**, orange compared to yellow), which is in qualitative agreement with reports of chylomicron remnants clear after approximately 6- to 8-hours after eating (23).

In conclusion of the meal intervention predictions, the model indicates qualitative agreement to many clinical observations in circulating CM levels and provides a small-scale mathematical model capable of simulating CM postprandial dynamics.

## Discussion

We have constructed a mathematical model describing plasma CM appearance (**Fig. 1A**) and evaluated its ability to describe data and utility in several steps (**Fig. 1B**). First, we found that the model could sufficiently describe six different clinical studies individually (**Fig. S1**; confirmed using individual χ^2^-test). Second, we found that the most important parameters to describe group variability (study and intervention) were *kL1* (governing enterocyte transport of lipid) and *kBreakdown* (CM clearance from plasma) (**Figs. S2-S3**). By allowing freedom in these two parameters and restricting variability in all other parameters, the model was able to simultaneously fit all estimation data (**Fig. 2**), confirmed by a χ^2^-test (90.1 < 148.8 = Tχ^2^). A relationship between the intervention-specific *kL1* and *kBreakdown* to study-specific BMI was found using linear regression (**Fig. 3A**). Third, the established BMI-specific calibration was used to predict new independent validation data (**Fig. 3B**). Finally, we investigated how the model could be used to predict postprandial chylomicron dynamics of various meal interventions for people with various BMIs (**Fig. 4**). In summary, we have constructed a small-scale mathematical model of CM appearance and the SME, providing a foundation for creating and extending models describing postprandial lipid metabolism.

The model’s ability to predict SME under various conditions needs to be further evaluated. When the model was trained using only the Barrows et al. (9) dataset, the model could produce a sharp second peak (**Fig. S1L**), but when trained simultaneously to all studies, the peak is less prominent (**Fig. 2B**). This is mainly because the SME peak magnitude varies from study to study, e.g., peak in Vors et al. (13) compared to Barrows et al. (9) (**Fig. 2L** compared to **Fig. 2B**). These discrepancies could be due to several factors not accounted for in the mathematical model and in many cases are not reported in studies herein, such as differences in how CM was measured, the form of consumed fats (oil, emulsion, or spread), or interpopulation variability not captured by BMI (e.g., insulin sensitivity). While BMI and insulin sensitivity have been reported to correlate significantly (24), we found weak agreement with the insulin resistant group in the Hodson et al. study despite using the BMI-dependent calibration (**Fig. 3E**). We advise future works to implement insulin sensitivity in their model predictions to potentially overcome this shortcoming. Despite these listed model limitations and potential future works, the mechanisms incorporated in our small-scale model are sufficient to describe differences in the SME (**Figs. 2-4**; as confirmed by individual χ^2^-test and qualitative agreement to literature).

There are several biological mechanisms that were omitted from the presented model. We do not include the production of CMs from plasma-derived *non-esterified fatty acids* (NEFA) and glycerol, nor the conversion of dietary nor plasma derived carbohydrate conversion into CM. Furthermore, there are other aspects to expand upon: gastric emptying crosstalk interactions between different macronutrients and meal volumes, and the fact that different fatty acids may have different metabolic rates. We have recently published an ethanol rate of appearance model, with a gastric emptying module (25), and future work could involve integration of such gastric emptying mechanisms to better describe lipid absorption, or interaction with other foods. Furthermore, as mentioned earlier, in a recent work by Sosa et al. (11) a comprehensive model is presented, which includes several of these pathways. Thus, in future work, we will evaluate if one could make use of the description of more detailed mechanisms described in the Sosa et al. model, to achieve a better simultaneous agreement to all data.

The relationship between covariates such as BMI and HOMA-IR in relation to their effect on chylomicron dynamics needs to be evaluated. There exists a known relationship between metabolic dysregulation and elevated levels of circulating TAGs, where mechanisms are e.g. poor clearance of CM-TAGs, or increased *Very Low Density Lipoprotein* (VLDL)-TAG secretion (26). The model analysis indicated the clearance of CM from plasma (*kBreakdown*) as significant for describing interstudy variability, and we show that this parameter has a tentative correlation with BMI (**Fig. 3A**). The perceived correlation with BMI and model parameters could be reasoned to be explained by the decreased clearance due to a change in insulin sensitivity. Yet, when we tested this correlation, we did not appear to find a similar correlation between the breakdown parameter and HOMA-IR (**Fig. S6**). A larger dataset with more covariate differences would be beneficial to understand these relationships further. Nevertheless, doing model personalization as presented herein could possibly be a strong approach if further developed, connecting rate parameters with individual anthropometrics, after the model training (not during).

There are several additional key aspects to consider in our study. The first aspect is regarding data availability. All data used for model estimation and validation is digitized data, thus the true data points might be slightly different from the ones we used. In many cases there was a lot of covariate data missing (such as BMI, age, and HOMA-IR), which excluded several studies from the analysis done (4,9). Furthermore, a delimitation of the study was not to include plasma TAG data, which further excluded a lot of available datasets. By including plasma TAG, the model complexity would also need to increase, for inclusion of various lipoproteins, i.e. VLDL. We have previously developed models for NEFA metabolism (27), lipolysis (28), glucose-insulin regulation (29,30), and multi-level and multi-timescale models for insulin resistance progression (31,32). An integration of all these models into a singular framework could be capable of describing mixed meal postprandial metabolism and therefore connecting more complex mechanisms of obesity progression and dieting. Herein, we present a step towards such a more complete mathematical model of human meal responses.

In conclusion, we have created a model for plasma CM appearance capable of describing dynamics related to the SME. The mechanism behind CM, such as person-specific variability and form of consumed type of dietary fats (oil, emulsion, or spread), is still not fully understood, and there seems to be large inter-patient, and inter-cohort variations in CM dynamics that will need further work to be elucidated. Nevertheless, our presented model is usable and could be integrated into any mathematical model for meal simulations, needing a description of CM appearance and postprandial lipid dynamics.

## Method

The code to reproduce all reported observations can be found in our GitHub repository (https://github.com/chrsi30/CMDT).

### Modelling approach and software

The model was formulated using ordinary differential equations (ODEs). The mathematical analysis, model simulation, model formulation and the model parameter estimation were all performed in MATLAB (2022a; The MathWorks, Natick, MA), using the systems biology toolbox (IQM) (33). The parameter estimation was performed using the extended scatter search (ESS) optimization algorithm from the MEIGO toolbox (34). The quality of the resulting optimal simulations was evaluated using a *χ*^2^-test (35). Lastly, the parameter and model uncertainty analysis was done using Markov-Chain Monte Carlo sampling implemented in the PESTO toolbox (36).

### Parameter estimation

Parameter estimation was done by quantifying the model performance, using the model output ŷ to calculate a weighted least squares cost function (**Eqs. 1-2**).

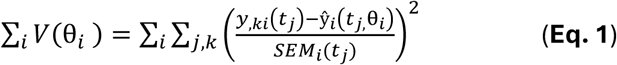

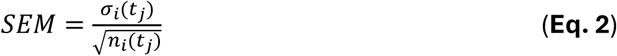

where θ is the model parameters; *y_i_*(*t_j_*) is the measured data from a study *i*, at time point *j and* from one type of measure *k*; ŷ*_i_*(*t_j_*,*q*) is the simulation value for a given experiment setup *i* and time point *j*, and SEM is the standard error of the mean, which is the sample standard deviation,*σ_i_*(*t_j_*) divided with the square root of the number of repeats, *n_i_*(*t_j_*) at each time point. The value of the cost function, *V*(θ), is then minimized by tuning the values of the parameters.

To evaluate the model, a *χ*^2^-test was performed for the size of the residuals, with the null hypothesis that the experimental data have been generated by the model. The test is performed under the assumption that the experimental noise is additive and normally distributed (35). In practice, the cost function value was compared to a *χ*^2^-test statistic, *T°*, with a cumulative density function (**Eq. 3**).

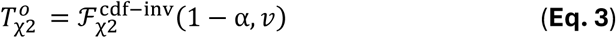

where 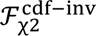 is the inverse density function; and α is the significance level (α = 0.05 was used) and *v* is the degrees of freedom, equal to the number of data points in the training dataset. In practice, the model is rejected if the model cost is larger than the *χ*^2^-threshold (*T°*).

### Model description – the chylomicron model

The ODE model was developed by the iterative process of physiologically based data-driven hypothesis testing. The capability of several different candidate model structures was tested using the Barrows et al. dataset. The candidates were evaluated based on their ability to accurately describe the experimental data using a χ^2^-test, with special emphasis on the ability to reproduce the sharp second meal peak. Furthermore, the candidate structure complexity was also evaluated based on the Akaike information criterion (AIC). The results of this evaluation can be seen in the supplementary material (**Table S2**). The chosen model (‘M9’) had the lowest χ^2^-test statistic, and it could best describe the sharp second meal peak observed in data. This section will detail the model equations.

The model has a total of seven model states and five model parameters (**Fig. 1A, right**). The first state represents TAG that has entered the stomach and is moving towards the intestine and has notation LS (**Eq. 4**).

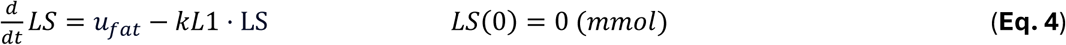

where; *u_fat_* is the model input, the mass of triglycerides from the meal, expressed as mmol (converted from grams using a TAG molar mass of 845mol/g), and *kL1* (min^-1^) is the rate parameter for the absorption and transfer of lipids into and within the enterocytes. TAG is then moved through the Enterocytes and prepared for storage or plasma release. This part is represented by the three model states: L1 (**Eq. 5**), L2 (**Eq. 6**), and L3 (**Eq. 7**).

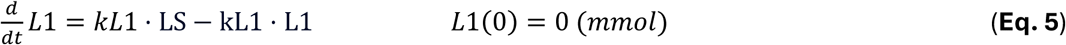

Where *kL1* again is the rate of transfer.

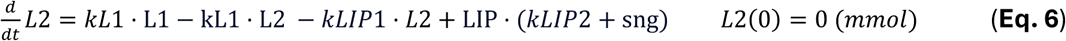

where *kL1* is reused from (Eq. 5) to describe the transfer rate moving into the next enterocyte state. In (Eq. 6) enterocyte lipid storage is introduced, which is the movement of TAG into lipid droplets represented by the state LIP. The transfer into and from the LIP state is represented by the parameters kLIP1 (min^-1^) and kLIP2 (min^-1^) respectively. Here we also introduce the phenomenological effect of the second meal, denoted *sng*, which affects the efflux from the LIP state.

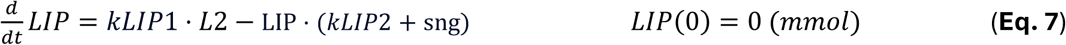

The sng effect is a model state which has the following derivative (**Eq. 8**):

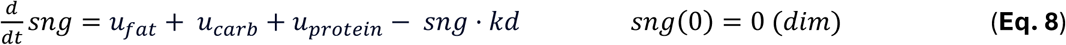

where *u_carb_* and *u_protein_* are the carbohydrate and protein portions of the meal. The last enterocyte state, L3, has the following derivative (**Eq. 9**):

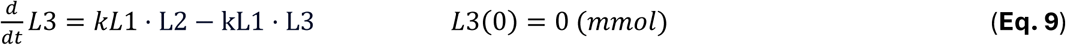

where *kL1* is reused from (Eqs. 5 and 6) as the rate of TAG-chylomicron excretion into plasma. The plasma TAG-chylomicron state is denoted *Chp* and has the following derivative (**Eq. 10**):

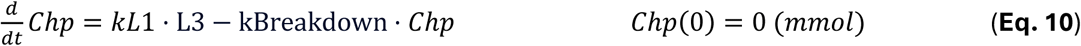

Where, *kBreakdown* (min^-1^) is the breakdown and usage of TAG-chylomicrons in the body. To obtain the concertation in plasma the *Chp* state is divided into with plasma volume, calculated via the Nadler formula, based on sex, height and weight (37).

### Data curation

Data was gathered from a total of seven published studies (4,5,9,12,13,16–18). The estimation data includes six of these studies and was comprised of a total of 10 different cohorts (**Table S1**), and various interventions. The interventions that were investigated in each study are described in the supplementary material (**Table S3**). Inclusion criteria for data were that TAG chylomicrons concentration should be measured in connection to a meal, and detailed information about the meal, and any prior meal should be detailed. All data was digitized using the WebPlotDigitzer.

## Author contributions

CS: Conceptualization, Methodology, Software, Formal analysis, Investigation, Data Curation, Writing – Original Draft, Visualization. OS: Conceptualization, Methodology, Software, Formal analysis, Investigation, Writing – Original Draft, Visualization. HP: Conceptualization, Methodology, Writing - Review & Editing. KT: Conceptualization, Methodology, Writing - Review & Editing. WL: Supervision, Methodology, Writing – Original Draft. KS: Supervision Writing - Review & Editing, Funding Acquisition, Writing - Review & Editing. EN: Supervision Investigation, Conceptualization, Methodology, Supervision, Funding Acquisition, Writing - Review & Editing. GC: Supervision Funding Acquisition, Project Administration, Investigation, Conceptualization, Methodology, Supervision, Writing – Original Draft.

## Supplementary material

**Figure S1.**
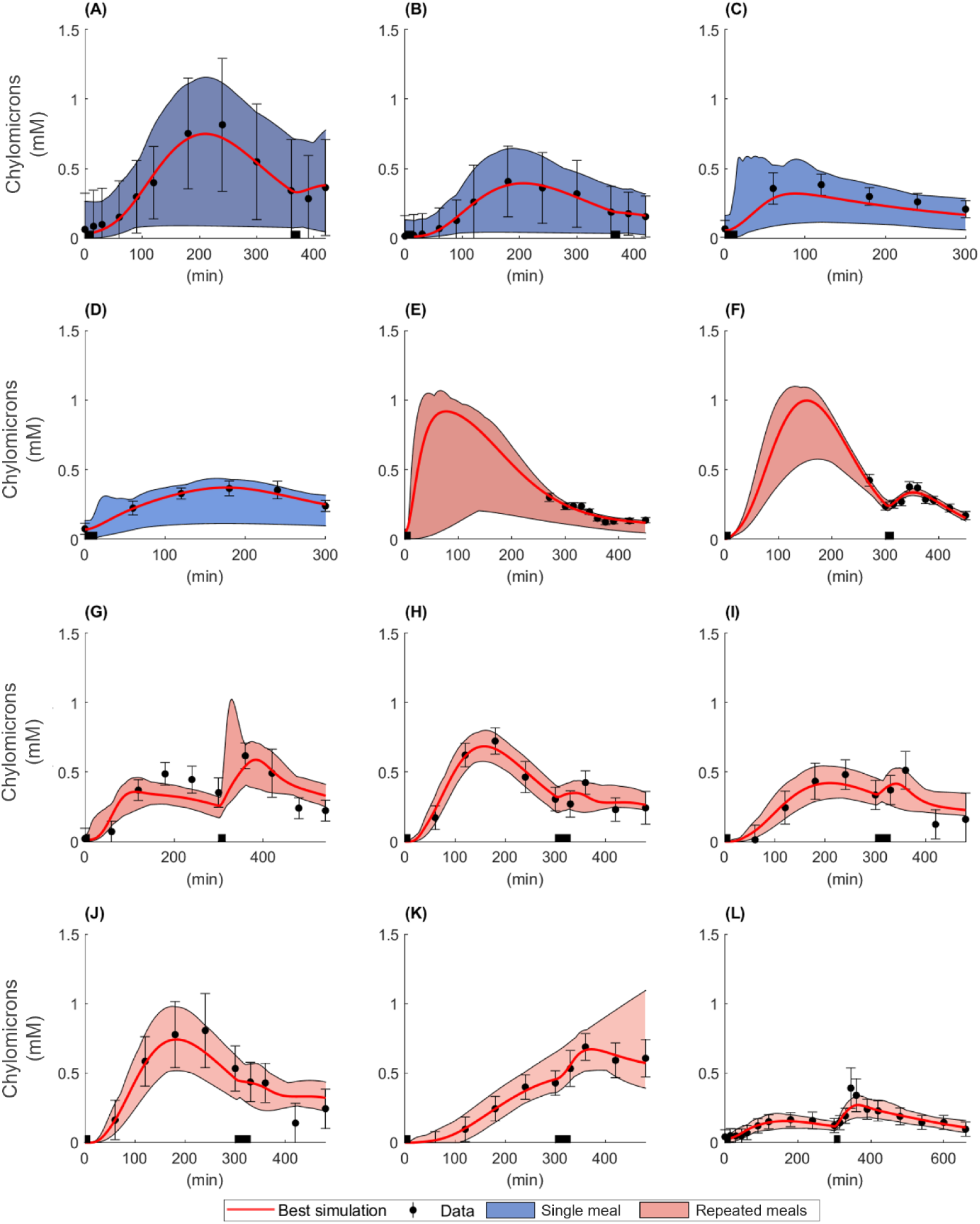
Model agreement to estimation data fitted to each study individually. For all panels: area indicates model uncertainty, errorbars represent data, lines represent model best agreement with data, and thick black bar on x-axis indicate meal consumption. **(A)** Model agreement for the Pramfalk et al., (12) male dataset. **(B)** Pramfalk et al., (12) female dataset **(C)** Milan et al., (16) young **(D)** Milan et al., (16) old **(E)** Evans et al., (4) low-fat **(f)** Evans et al. (4) water **(G)** Heath et al. (17) healthy **(H)** Vors et al. (13) normal spread **(I)** Vors et al. (13) normal emulsion **(J)** Vors et al. (13) obese spread. **(K)** Vors et al. (13) obese emulsion **(L)** Barrows et al. (9) two identical meals.

**Figure S2.**
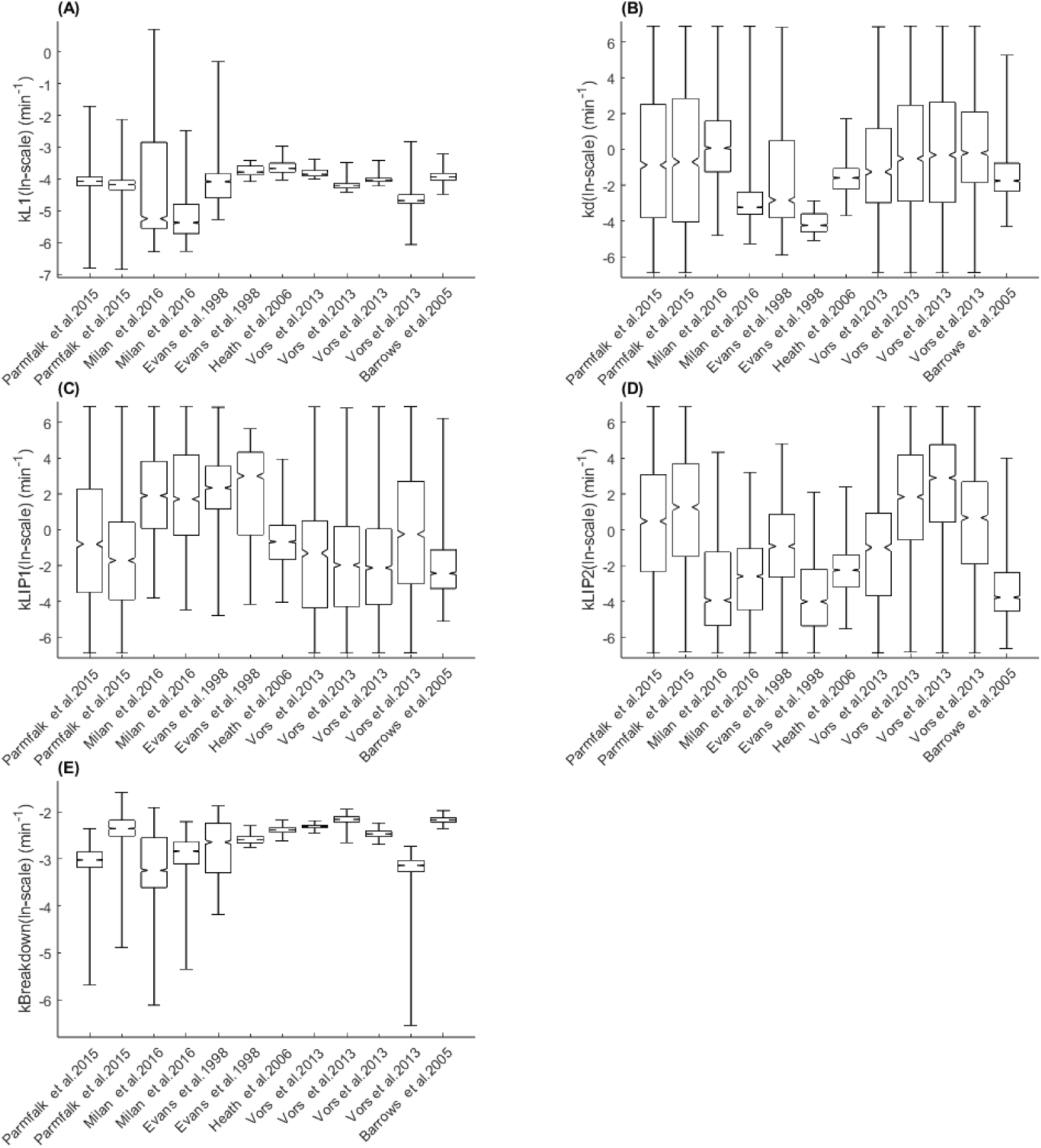
Parameter coefficients when model is trained to all estimation data individually. Boxplots showing the parameter distribution for model parameters. **(A)** kL1. **(B)** kd. **(C)** kLIP1. **(D)** kLIP2. **(E)** kBreakdown.

**Figure S3.**
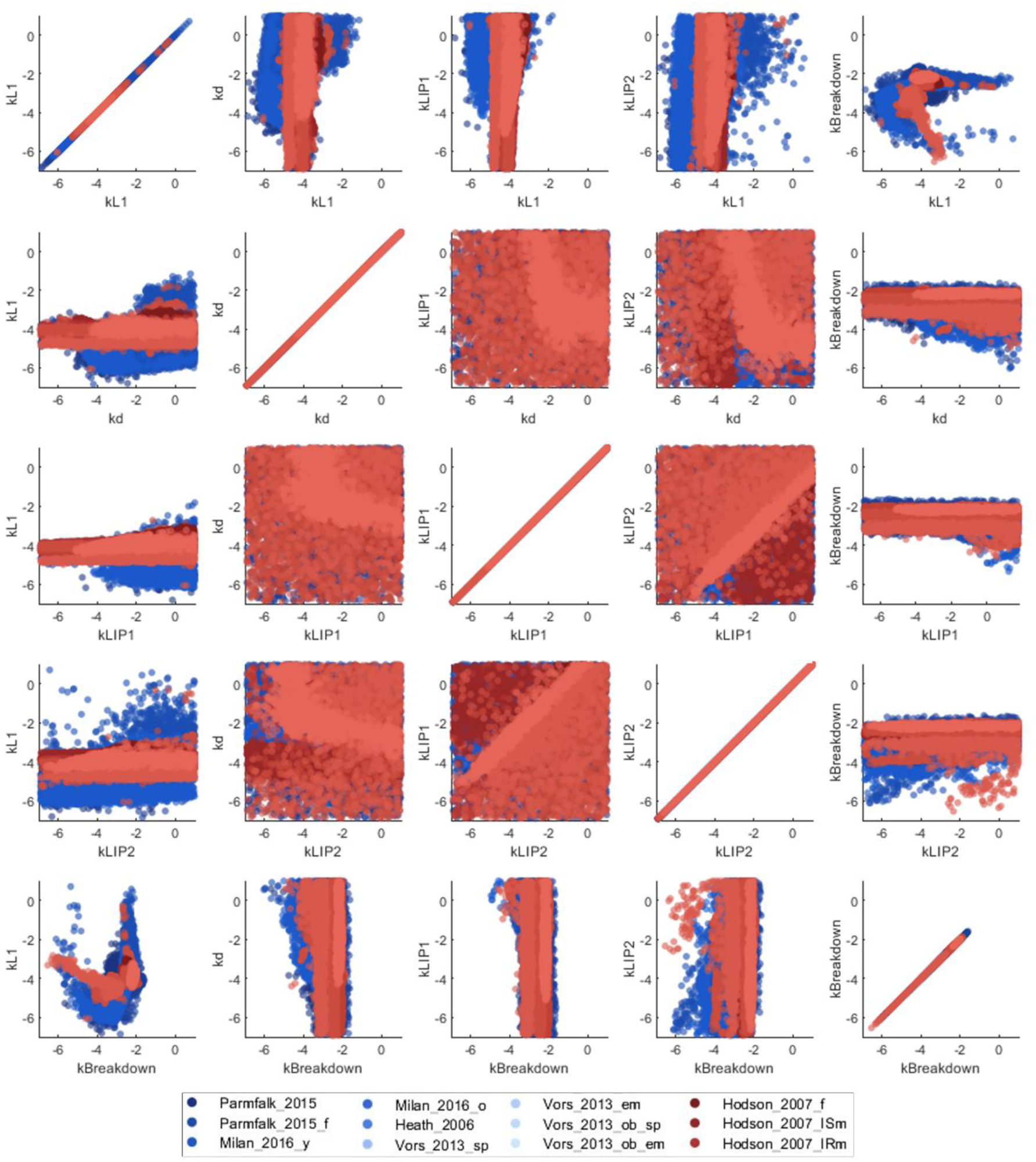
2D parameter profiles after sampling. This figure shows the relationships between all parameters during the individual agreement to data. The different colors represent the different studies (**Table S1**), with the blue colored dots being parameters estimated to single-meal studies, and red dots for studies with sequential meals.

**Figure S4.**
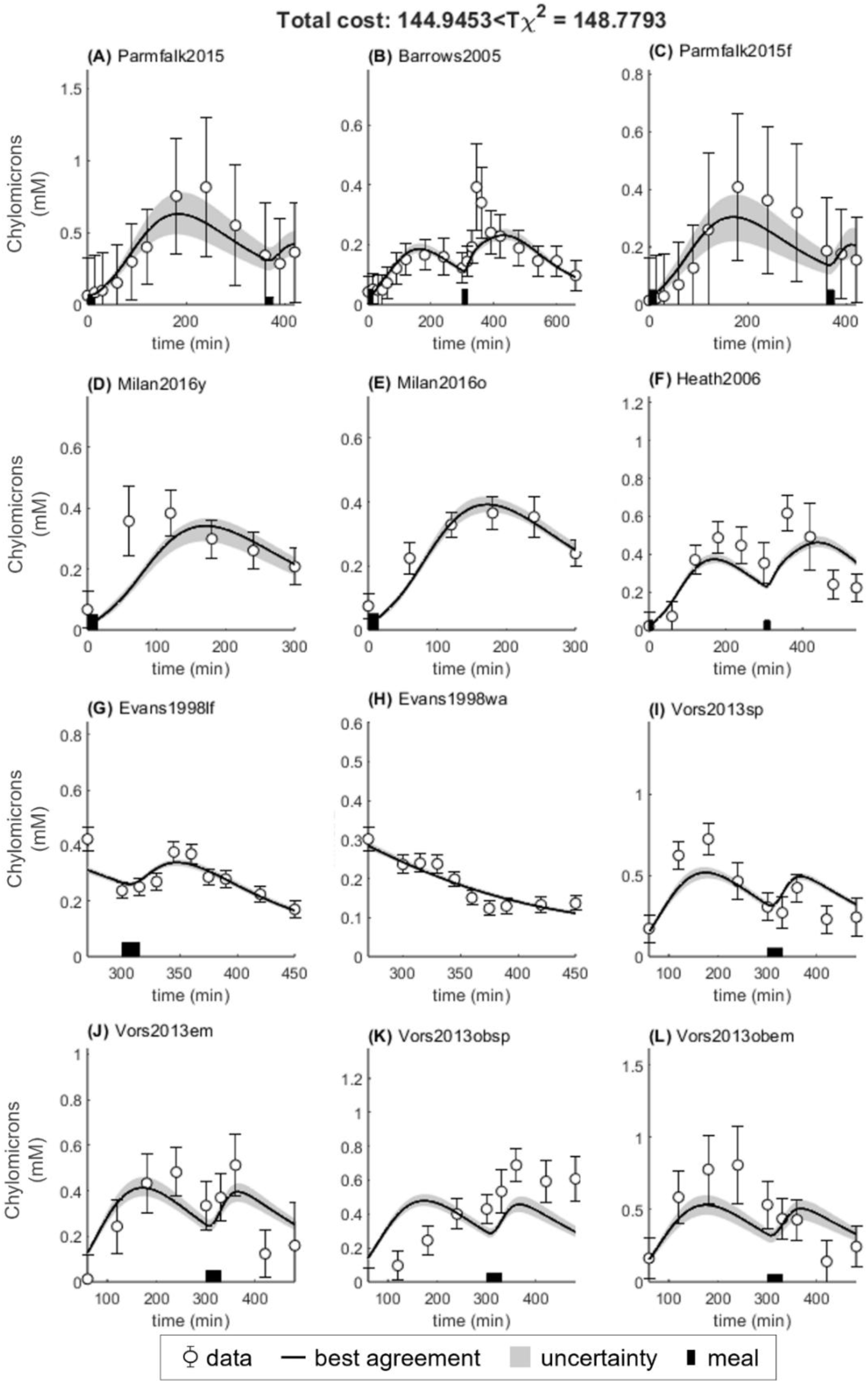
Model simultaneous agreement to all estimation data with allowed variability only in parameter kBreakdown and not clearance. For all panels: area indicates model uncertainty, errorbars represent data, lines represent model best agreement with data, and thick black bar on x-axis indicate meal consumption. **(A)** Pramfalk et al., (12) males **(B)** Barrows et al., (9) healthy. **(C)** Pramfalk et al., (12) females. **(D)** Milan et al., (16) young, **(E)** Milan et al., (16) old. **(F)** Heath et al., (17) healthy. **(G)** Evans et al., low-fat (4). **(H)** Evans et al., water (4). **(I)** Vors et al., normal spread. **(J)** Vors et al., normal emulsion (13). **(K)** Vors et al., obese spread (13). **(L)** Vors et al., obese emulsion (13).

**Figure S5.**
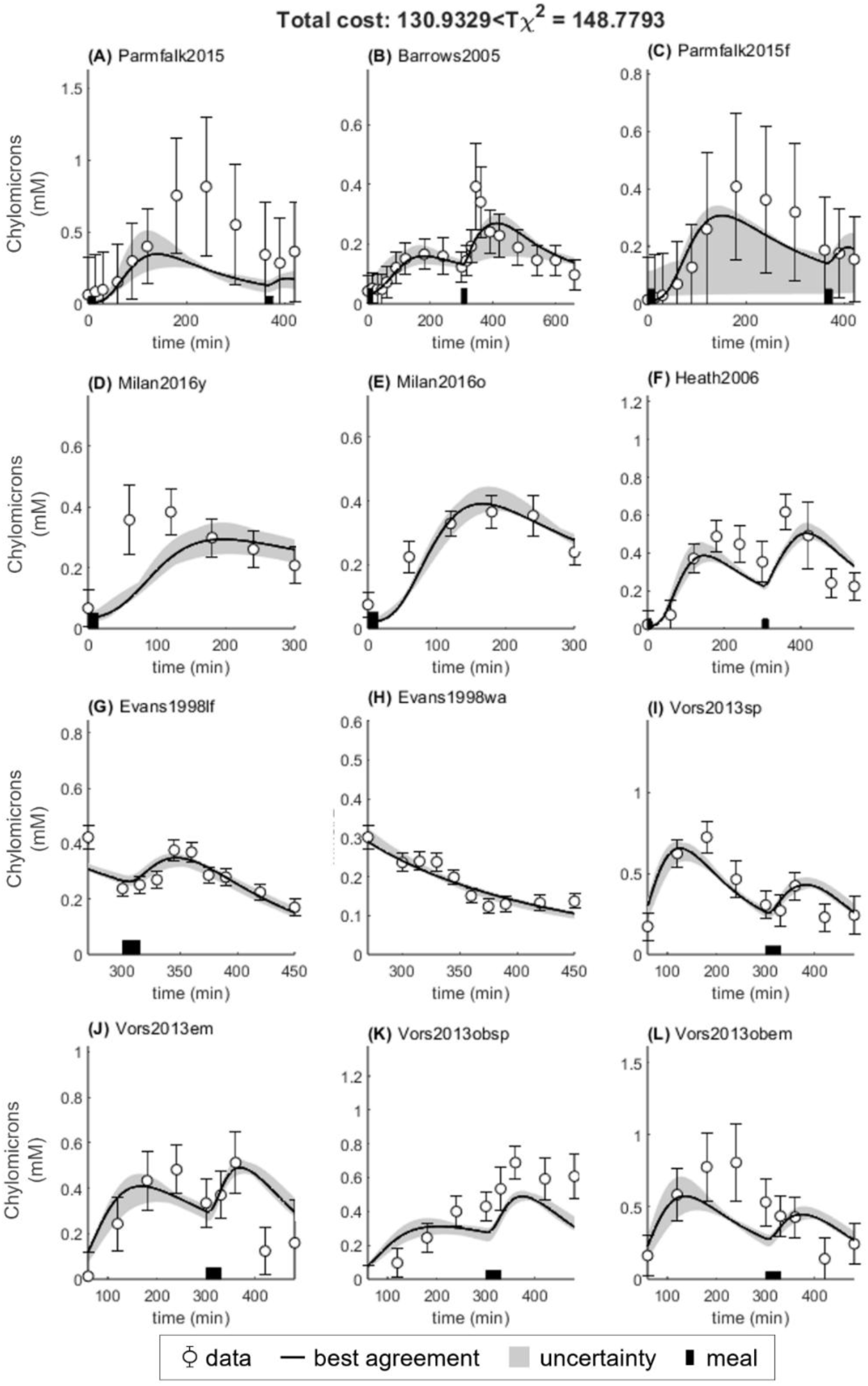
Model simultaneous agreement to all estimation data with allowed variability only in clearance and not kBreakdown. For all panels: area indicates model uncertainty, errorbars represent data, lines represent model best agreement with data, and thick black bar on x-axis indicate meal consumption. **(A)** Pramfalk et al., (12) males **(B)** Barrows et al., (9) healthy. **(C)** Pramfalk et al., (12) females. **(D)** Milan et al., (16) young, **(E)** Milan et al., (16) old. **(F)** Heath et al., (17) healthy. **(G)** Evans et al., (4) low-fat. **(H)** Evans et al., (4) water. **(I)** Vors et al., (13) normal spread. **(J)** Vors et al., (13) normal emulsion. **(K)** Vors et al., (13) obese spread. **(L)** Vors et al., (13) obese emulsion.

**Figure S6.**
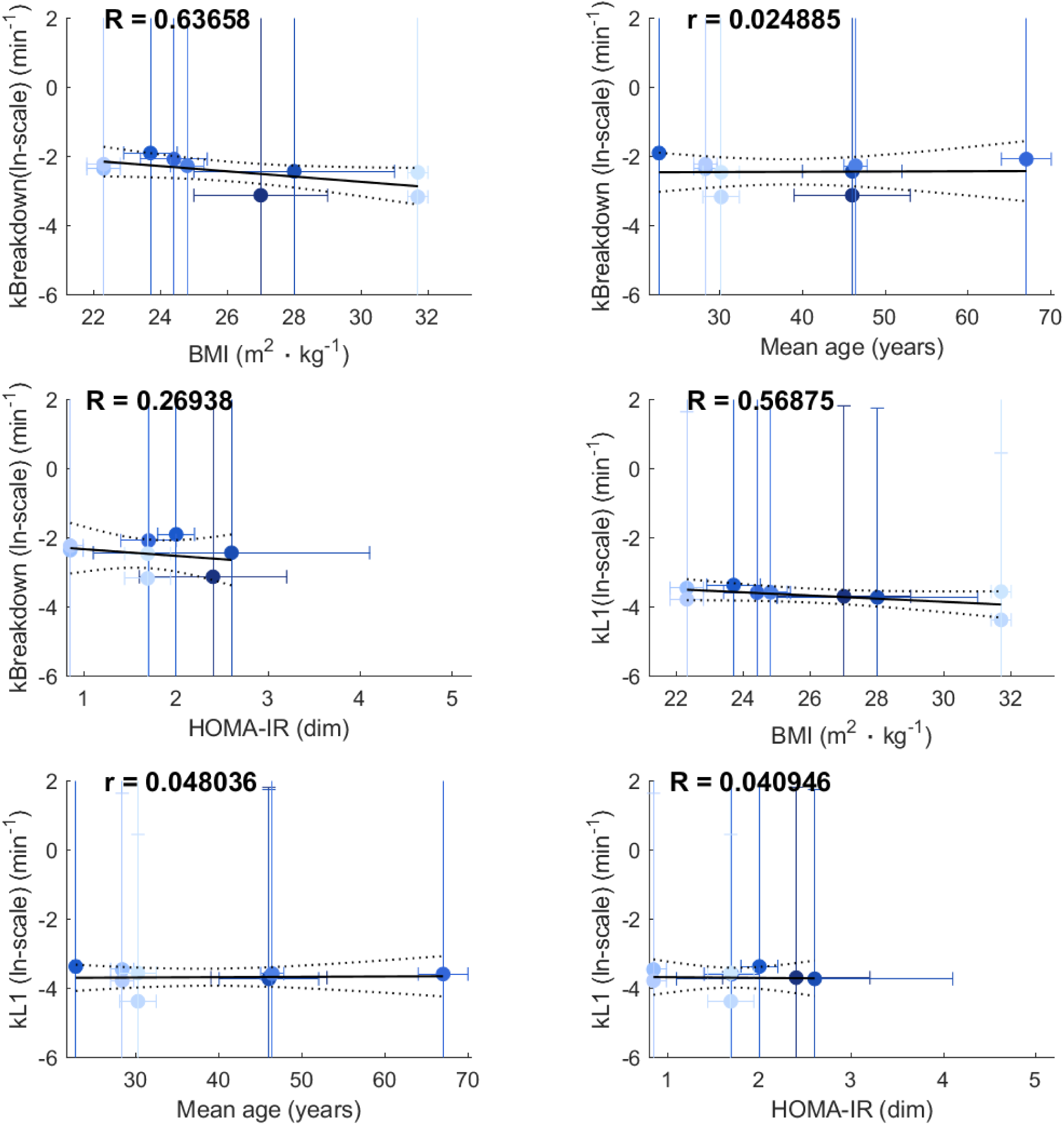
Regression analysis between calibration parameters kBreakdown and *kL1* to possible covariates. (A) *kBreakdown* vs BMI. (B) *kBreakdown* vs age. (C) *kBreakdown* vs HOMA-IR. (D) *kL1* vs BMI. (E) *kL1* vs age. (F) *kL1* vs HOMA-IR.

**Table S1.** Table giving a brief description of the anthropometric data and clinical parameter characterizing the study cohort used for training and validation data.

| Study name | Cohort | Age | BMI | HOMA-IR |
| --- | --- | --- | --- | --- |
| **Barrows et al. 2005** | Healthy (M) (n=6) | 20–55 | - | - |
| **Evans et al. 1998** | Healthy (M/F) (n=6) | 21–44 | 20.8–27.2 | - |
| **Milan et al. 2016** | Healthy (M/F) (n = 15) | 67.3±1.5 | 24.4±1.0 | 1.7±0.3 |
| **Milan et al. 2016** | Healthy (M/F) (n = 15) | 22.7±0.4 | 23.7±0.8 | 2.0±0.2 |
| **Vors et al. 2013** | Healthy (n=9) | 28.3±1.4 | 22.3±0.5 | 0.85±0.14 |
| **Vors et al. 2013** | Obese (n=9) | 30.2±2.2 | 31.7±0.3 | 1.69±0.25 |
| **Heath et al. 2006** | Healthy (M) (n=8). | 46.4±1.4 | 24.8±0.5 | - |
| **Pramfalk et al. 2015** | Healthy (F) (n=11). | 46±6 | 28±3 | 2.6 ±1.5 |
| **Pramfalk et al. 2015** | Healthy (M) (n=11). | 46±7 | 27±2 | 2.4±0.8 |
| **Hodson et al.** | Healthy(F) (n=11). | 32 (23–64) | 22 (22–27) | 0.9 (0.4–1.3) |
| **Hodson et al.** | Insulin sensitive (M) (n=11) | 42 (23–54) | 26 (21–31) | 1.1 (0.9–1.4) |
| **Hodson et al.** | Insulin resistant (M) (n=11) | 42 (22–45) | 25 (22–35) | 1.7 (1.5–3.5) |

**Table S2.** We evaluated several candidate model structures before choosing ‘M9’ as the final model structure that could meet our criteria; agreement to the Barrows et al. data with prominent SME.

| Model Name | Description | Params | States | Training cost (V(θ) (Barrows et al)) |
| --- | --- | --- | --- | --- |
| **M1** | State for meal effect on LIP outflux, with chylomicron dependency. 3 transfer states. Two different rate parameters for transfer states. | **6**<br>(*kL1,kL2,kd,kLIP1,kLIP2,kBeakdown*) | 7 | 4.5 |
| **M2** | But the state for meal effect was variable. (**Eq. 8**) | **6**(*kL1,kL2,kd,kLIP1,kLIP2,kBreakdown*) | 6 | 6.2 |
| **M3** | But there is a single parameter for governing all transfer states. (**Eq.4,5,6 and 9**) | **5**(*kL1,kd,kLIP1,kLIP2,kBreakdown*) | 7 | 4.48 |
| **M4** | Using only one parameter for influx and outflux from LIP state. (**Eq. 6-7**) | **4**(*kL1,kd,kLIP1,kBreakdown*) | 7 | 22.14 |
| **M5** | Removed on delay state. | **5**(*kL1,kd,kLIP1,kLIP2,kBreakdown*) | 6 | 6.47 |
| **M6** | Removed two delay states | **5**(*kL1,kd,kLIP1,kLIP2,kBreakdown*) | 5 | 9.42 |
| **M7** | But the state for meal effect was variable. (**Eq. 8**) | **4**(*kL1,kLIP1,kLIP2,kBreakdown*) | 6 | 6.3 |
| **M8** | Removed chylo-effect on LIP outflux. | **5**(*kL1,kd,kLIP1,kLIP2,kBreakdown*) | 7 | 2.6 |
| **M9** | Using only one parameter for influx and outflux from LIP state. (**Eq. 6-7**) | **4**(*kL1,kd,kLIP1,kBreakdown*) | 7 | 8.9 |

**Table S3.** Table giving a brief description of the study intervention used for the model estimation and validation data.

| Study name | Description of meal | Fat content (g) |
| --- | --- | --- |
| **Pramfalk *et al.* 2015** | Overnight fasting followed by two meals. 40g Rice Krispies (Kellogg), 200g skimmed milk, and a chocolate milkshake containing 40g of olive oil. The second meal was 75g glucose. | Estimated.<br>Breakfast: ~42.4g |
| **Barrows *et al.* 2005** | Overnight fasting followed by two meals. The total energy provided by the liquid formula was $7,260 \pm 1,346$ kJ, with a macronutrient profile of 54.1% of energy from carbohydrates, 32.1% from fat, and 14.1% from protein. Liquid meals. Pasteurized egg yolk, heavy whipping cream, and vegetable oil. | Estimated ~36g.<br>7260kJ.<br>$0.2390 \text{ kcal} \cdot \text{kJ}^{-1}$<br>9kcal/g |
| **Evans *et al.* 1998** | Overnight fasting followed by two meals. High-fat breakfast and lunch. The lunch was either low-fat or high fat. Breakfast: Safflower oil (55g), egg yolk, egg white, bread, jam. High-fat lunch: full-cream milk, cornflakes, full-fat ice cream, and double cream. Low-fat lunch: skimmed milk, cornflakes, and fat-free ice cream. | Given.<br>Breakfast: 62.9g<br>High-fat lunch: 56.9g<br>Low-fat lunch: 0.9g |
| **Heath *et al.* 2006** | Overnight fasting followed by two meals. Breakfast, consisting of 40g of Rice Krispies (Kellogg), a banana, and chocolate milkshake containing 50g of fat. The fat content of lunch was also 50g. It consisted of a cheese sandwich and a second warm chocolate milkshake. | Given.<br>Breakfast: 51.9g<br>Lunch: 49.6g |
| **Milan *et al.* 2016** | Overnight fasting followed by a single meal. Mixed meal breakfast | Given.<br>Breakfast: 62.2g |
| **Vors *et al.* 2013** | Overnight fasting followed by two meals. Breakfast either as fat spread or fat emulsion. Lunch was a mixed meal. Both consisted of bread (50g), skimmed milk, and anhydrous milk fat (40g) containing 600mg tracers—either spread on bread or emulsified in skim milk. A second meal was served 5 h after breakfast. The meal contained pasta (200g), turkey (100g), butter (10g), olive oil (10g), bread (50g), and stewed fruit (100g), which provided 713 kcal (2985 kJ) with 29%, 51%, and 20% of energy as lipids (22.7g), carbohydrates (91.5g), and proteins (35.7g), respectively. | Given.<br>Breakfast: 41.4g<br>Lunch: 22.7g |

## References

1. Bergman RN, Ider YZ, Bowden CR, Cobelli C. Quantitative estimation of insulin sensitivity. American Journal of Physiology-Endocrinology and Metabolism. 1979 Jun 1;236(6):E667. doi:10.1152/ajpendo.1979.236.6.E667

2. Dalla Man C, Rizza RA, Cobelli C. Meal simulation model of the glucose-insulin system. IEEE Trans Biomed Eng. 2007 Oct;54(10):1740–9. doi:10.1109/TBME.2007.893506

3. Gugliucci A. The chylomicron saga: time to focus on postprandial metabolism. Front Endocrinol (Lausanne). 2023;14:1322869. doi:10.3389/fendo.2023.1322869 PubMed PMID: 38303975; PubMed Central PMCID: PMC10830840.

4. Evans K, Kuusela PJ, Cruz ML, Wilhelmova I, Fielding BA, Frayn KN. Rapid chylomicron appearance following sequential meals: effects of second meal composition. British Journal of Nutrition. 1998 May;79(5):425–9. doi:10.1079/BJN19980072

5. Fielding B, Callow J, Owen R, Samra J, Matthews D, Frayn K. Postprandial lipemia: the origin of an early peak studied by specific dietary fatty acid intake during sequential meals. The American Journal of Clinical Nutrition. 1996 Jan 1;63(1):36–41. doi:10.1093/ajcn/63.1.36

6. Lambert JE, Parks EJ. Postprandial metabolism of meal triglyceride in humans,. Biochim Biophys Acta. 2012 May;1821(5):721–6. doi:10.1016/j.bbalip.2012.01.006 PubMed PMID: 22281699; PubMed Central PMCID: PMC3588585.

7. Jackson KG, Robertson MD, Fielding BA, Frayn KN, Williams CM. Second meal effect: modified sham feeding does not provoke the release of stored triacylglycerol from a previous high-fat meal. British Journal of Nutrition. 2001 Feb;85(2):149–56. doi:10.1079/BJN2000226

8. Jackson KG, Robertson MD, Fielding BA, Frayn KN, Williams CM. Olive oil increases the number of triacylglycerol-rich chylomicron particles compared with other oils: an effect retained when a second standard meal is fed1,2,3,4. The American Journal of Clinical Nutrition. 2002 Nov 1;76(5):942–9. doi:10.1093/ajcn/76.5.942

9. Barrows BR, Parks EJ. Contributions of Different Fatty Acid Sources to Very Low-Density Lipoprotein-Triacylglycerol in the Fasted and Fed States. The Journal of Clinical Endocrinology & Metabolism. 2006 Apr 1;91(4):1446–52. doi:10.1210/jc.2005-1709

10. Mattes RD. Oral Exposure to Butter, but Not Fat Replacers Elevates Postprandial Triacylglycerol Concentration in Humans. The Journal of Nutrition. 2001 May 1;131(5):1491–6. doi:10.1093/jn/131.5.1491

11. Jacome-Sosa M, Hu Q, Manrique-Acevedo CM, Phair RD, Parks EJ. Human intestinal lipid storage through sequential meals reveals faster dinner appearance is associated with hyperlipidemia. JCI Insight. 6(15):e148378. doi:10.1172/jci.insight.148378 PubMed PMID: 34369385; PubMed Central PMCID: PMC8489663.

12. Pramfalk C, Pavlides M, Banerjee R, McNeil CA, Neubauer S, Karpe F, et al. Sex-Specific Differences in Hepatic Fat Oxidation and Synthesis May Explain the Higher Propensity for NAFLD in Men. The Journal of Clinical Endocrinology & Metabolism. 2015 Dec 1;100(12):4425–33. doi:10.1210/jc.2015-2649

13. Vors C, Pineau G, Gabert L, Drai J, Louche-Pélissier C, Defoort C, et al. Modulating absorption and postprandial handling of dietary fatty acids by structuring fat in the meal: a randomized crossover clinical trial1 2 3. The American Journal of Clinical Nutrition. 2013 Jan 1;97(1):23–36. doi:10.3945/ajcn.112.043976

14. Leohr J, Heathman M, Kjellsson MC. Semi-physiological model of postprandial triglyceride response in lean, obese and very obese individuals after a high-fat meal. Diabetes, Obesity and Metabolism. 2018 Mar;20(3):660–6. doi:10.1111/dom.13138

15. O’Donovan SD, Erdős B, Jacobs DM, Wanders AJ, Thomas EL, Bell JD, et al. Quantifying the contribution of triglycerides to metabolic resilience through the mixed meal model. iScience. 2022 Nov 18;25(11):105206. doi:10.1016/j.isci.2022.105206 PubMed PMID: 36281448; PubMed Central PMCID: PMC9587016.

16. Milan AM, Nuora A, Pundir S, Pileggi CA, Markworth JF, Linderborg KM, et al. Older adults have an altered chylomicron response to a high-fat meal. British Journal of Nutrition. 2016 Mar;115(5):791–9. doi:10.1017/S000711451500505X

17. Heath RB, Karpe F, Milne RW, Burdge GC, Wootton SA, Frayn KN. Dietary fatty acids make a rapid and substantial contribution to VLDL-triacylglycerol in the fed state. American Journal of Physiology-Endocrinology and Metabolism. 2007 Mar;292(3):E732–9. doi:10.1152/ajpendo.00409.2006

18. Hodson L, Bickerton AST, McQuaid SE, Roberts R, Karpe F, Frayn KN, et al. The Contribution of Splanchnic Fat to VLDL Triglyceride Is Greater in Insulin-Resistant Than Insulin-Sensitive Men and Women : Studies in the Postprandial State. Diabetes. 2007 Oct 1;56(10):2433–41. doi:10.2337/db07-0654

19. Wong ATY, Chan DC, Pang J, Watts GF, Barrett PHR. Plasma apolipoprotein B-48 transport in obese men: a new tracer kinetic study in the postprandial state. J Clin Endocrinol Metab. 2014 Jan;99(1):E122–126. doi:10.1210/jc.2013-2477 PubMed PMID: 24203058.

20. Jegatheesan P, Seyssel K, Stefanoni N, Rey V, Schneiter P, Giusti V, et al. Effects of gastric bypass surgery on postprandial gut and systemic lipid handling. Clinical Nutrition ESPEN. 2020 Feb 1;35:95–102. doi:10.1016/j.clnesp.2019.11.002

21. Björnson E, Packard CJ, Adiels M, Andersson L, Matikainen N, Söderlund S, et al. Apolipoprotein B48 metabolism in chylomicrons and very low-density lipoproteins and its role in triglyceride transport in normo- and hypertriglyceridemic human subjects. J Intern Med. 2020 Oct;288(4):422–38. doi:10.1111/joim.13017 PubMed PMID: 31846520.

22. Evans K, Kuusela PJ, Cruz ML, Wilhelmova I, Fielding BA, Frayn KN. Rapid chylomicron appearance following sequential meals: effects of second meal composition. Br J Nutr. 1998 May;79(5):425–9. doi:10.1079/bjn19980072 PubMed PMID: 9682661.

23. Keirns BH, Sciarrillo CM, Koemel NA, Emerson SR. Fasting, non-fasting and postprandial triglycerides for screening cardiometabolic risk. J Nutr Sci. 2021 Sep 14;10:e75. doi:10.1017/jns.2021.73 PubMed PMID: 34589207; PubMed Central PMCID: PMC8453457.

24. Ferrannini E, Natali A, Bell P, Cavallo-Perin P, Lalic N, Mingrone G. Insulin resistance and hypersecretion in obesity. J Clin Invest. 1997 Sep 1;100(5):1166–73. doi:10.1172/JCI119628 PubMed PMID: 9303923.

25. Podéus H, Simonsson C, Nasr P, Ekstedt M, Kechagias S, Lundberg P, et al. A physiologically-based digital twin for alcohol consumption—predicting real-life drinking responses and long-term plasma PEth. npj Digital Medicine. 2024 May 3;7(1):112. doi:10.1038/s41746-024-01089-6

26. Ginsberg HN, Zhang YL, Hernandez-Ono A. Regulation of Plasma Triglycerides in Insulin Resistance and Diabetes. Archives of Medical Research. 2005 May 1; Current Trends in Diabetes 36(3):232–40. doi:10.1016/j.arcmed.2005.01.005

27. Sips FLP, Nyman E, Adiels M, Hilbers PAJ, Strålfors P, van Riel NAW, et al. Model-Based Quantification of the Systemic Interplay between Glucose and Fatty Acids in the Postprandial State. PLoS One. 2015;10(9):e0135665. doi:10.1371/journal.pone.0135665

28. Lövfors W, Ekström J, Jönsson C, Strålfors P, Cedersund G, Nyman E. A systems biology analysis of lipolysis and fatty acid release from adipocytes in vitro and from adipose tissue in vivo. PLoS One. 2021;16(12):e0261681. doi:10.1371/journal.pone.0261681 PubMed PMID: 34972146; PubMed Central PMCID: PMC8719686.

29. Silfvergren O, Simonsson C, Ekstedt M, Lundberg P, Gennemark P, Cedersund G. Digital twin predicting diet response before and after long-term fasting. PLoS Comput Biol. 2022 Sep 12;18(9):e1010469. doi:10.1371/journal.pcbi.1010469 PubMed PMID: 36094958; PubMed Central PMCID: PMC9499255.

30. Silfvergren O, Rigal S, Schimek K, Simonsson C, Kanebratt KP, Forschler F, et al. M4 drug discovery: human drug predictions from integrated pre-clinical insights exemplified with a GLP1-R agonist [Internet]. bioRxiv; 2025 [cited 2025 Dec 18]. p. 2025.11.03.686224. Available from: https://www.biorxiv.org/content/10.1101/2025.11.03.686224v1 doi:10.1101/2025.11.03.686224

31. Herrgårdh T, Hunter E, Tunedal K, Örman H, Amann J, Navarro FA, et al. Digital twins and hybrid modelling for simulation of physiological variables and stroke risk [Internet]. bioRxiv; 2022 [cited 2025 Sep 15]. p. 2022.03.25.485803. Available from: https://www.biorxiv.org/content/10.1101/2022.03.25.485803v1 doi:10.1101/2022.03.25.485803

32. Simonsson C, Lövfors W, Bergqvist N, Nyman E, Gennemark P, Stenkula KG, et al. A multi-scale *in silico* mouse model for diet-induced insulin resistance. Biochemical Engineering Journal. 2023 Feb 1;191:108798. doi:10.1016/j.bej.2022.108798

33. Schmidt H, Jirstrand M. Systems Biology Toolbox for MATLAB: a computational platform for research in systems biology. Bioinformatics. 2006 Feb 15;22(4):514–5. doi:10.1093/bioinformatics/bti799

34. Egea JA, Henriques D, Cokelaer T, Villaverde AF, MacNamara A, Danciu DP, et al. MEIGO: an open-source software suite based on metaheuristics for global optimization in systems biology and bioinformatics. BMC Bioinformatics. 2014 May 10;15(1):136. doi:10.1186/1471-2105-15-136

35. Cedersund G. Conclusions via unique predictions obtained despite unidentifiability--new definitions and a general method. FEBS J. 2012 Sep;279(18):3513–27. doi:10.1111/j.1742-4658.2012.08725.x

36. Stapor P, Weindl D, Ballnus B, Hug S, Loos C, Fiedler A, et al. PESTO: Parameter EStimation TOolbox. Bioinformatics. 2017;34(4):705–7. doi:10.1093/bioinformatics/btx676

37. Nadler SB, Hidalgo JU, Bloch T. Prediction of blood volume in normal human adults. Surgery. 1962 Feb 1;51(2):224–32. doi:10.5555/uri:pii:0039606062901666 PubMed PMID: 21936146.

